# Spent Mushroom Substrate Adsorbs Fusaric Acid to Protect the Cucumber Rhizosphere Microbiome and Alleviates Fusarium Wilt

**DOI:** 10.64898/2025.12.31.697147

**Authors:** Yuanhang Qu, Xiaomeng Liu, Lemeng Dong, Zhenhe Su, Harro Bouwmeester, Ping Ma, Qinggang Guo

## Abstract

Caused by *Fusarium oxysporum* f. sp. *cucumerinum*, cucumber Fusarium wilt threatens cucumber production worldwide. In addition to pesticides, using spent mushroom substrate (SMS) has a protective effect against Fusarium infection in cucumber, partly through effects on the microbiome. Fusaric acid (FA), a phytotoxic virulence factor secreted by Fusarium, disrupts both plant physiology and the rhizosphere microbial community. Herein, we investigated the relationship between SMS and this virulence factor. Bioassays demonstrated that SMS adsorbed FA both in culture supernatants and in soil. Adsorption behaviour conformed to the Langmuir–Freundlich isotherm model, with a maximum adsorption capacity of 52.72 μg/g. Kinetics followed a pseudo-second-order model, indicating physical and chemical adsorption. Scanning electron microscopy showed that SMS had a porous surface, facilitating FA capture. Fourier transform infrared spectroscopy and chemical blocking assays were used to identify secondary amide groups as the key binding sites. In a dual-compartment pot system designed to isolate adsorption from other direct effects, SMS treatment significantly reduced the FA concentration in the cucumber rhizosphere and mitigated FA-induced disease aggravation. This lowered the disease index by up to 25%. Bacterial 16S metabarcoding showed that FA disrupted the rhizosphere bacterial diversity and community structure. However, when FA was adsorbed by SMS, the microbial diversity and community stability were restored. FA reduced the abundance of sensitive taxa such as *Bacillus*. Meanwhile, SMS-adsorbed FA preserved its relative abundance, suggesting a selective protective effect for FA-sensitive rhizobacteria. These findings indicate that SMS protects cucumber against Fusarium by alleviating FA toxicity toward beneficial microbes. Through FA adsorption, SMS amendment stabilised the rhizosphere microbial community and reduced disease incidence. This highlights the potential of SMS as a sustainable and microbiome-friendly strategy for managing soil-borne diseases.

## 1. Introduction

Soil-borne diseases are a persistent and widespread challenge in agriculture, posing a considerable threat to sustainable and efficient crop production. Among them, the genus *Fusarium* comprises some of the most destructive plant pathogens, infecting a wide range of hosts and causing substantial yield losses worldwide (Brown et al., 2021). *Fusarium* species are well adapted to survive in soil for extended periods and employ multiple pathogenic strategies, including secreting various toxic secondary metabolites during infection. These metabolites disrupt plant physiological processes and weaken host defences, facilitating pathogen colonisation and disease progression (Jin et al., 2024).

Among these toxins, fusaric acid (FA) is a representative mycotoxin that has been identified as a key virulence factor in many *Fusarium*–host interactions. FA exacerbates disease development by inhibiting plant growth and inducing wilting and chlorosis through multiple mechanisms, such as disruption of plant hormone metabolism, induction of reactive oxygen species (ROS) accumulation, and damage to cell membrane integrity (Jin et al., 2024). In addition to its direct phytotoxicity, FA exhibits broad-spectrum antimicrobial activity, suppressing the growth and activity of beneficial rhizosphere bacteria through chemical–ecological mechanisms (Shao et al., 2025). Plant growth-promoting rhizobacteria (PGPR), such as *Bacillus* spp., are particularly sensitive, with growth rates reduced at FA concentrations as low as 10 µg/mL. Meanwhile, rhizosphere concentrations can reach up to 100 µg/mL during disease outbreaks (Notz et al., 2002). This diminishes the competitive capacity of PGPR in the rhizosphere ecosystem and indirectly facilitates the colonisation and spread of pathogenic fungi. FA inhibits the metabolic pathways of PGPR, suppressing antibiotic synthesis and biocontrol potential, further impairing their ecological function in the rhizosphere (Quecine et al., 2016). FA functions as both a virulence determinant and an ecological modulator that disrupts beneficial microbial communities and reshapes the rhizosphere microecology (Noman et al., 2025).

Reducing the bioavailability of FA in the rhizosphere is a novel strategy for the ecological management of soil-borne diseases. Adsorption technology is highly sustainable and has been widely applied for toxin removal in environmental remediation and food safety. Porous adsorbent materials, such as montmorillonite, biochar, and organically modified composites, can effectively remove mycotoxins through physical adsorption and chemical binding (Li et al., 2018; Fu et al., 2020). Adsorption-based strategies may offer ecological benefits in managing soil-borne diseases. Biochar could reduce tomato Fusarium wilt by adsorbing cell wall-degrading enzymes and toxins secreted by *Fusarium oxysporum* (Jaiswal et al., 2018). These findings highlight the potential of adsorption-based approaches as a promising direction for the ecological management of soil-borne diseases.

As a byproduct of edible mushroom cultivation, SMS is a waste rich in organic matter, cellulose, and residual fungal mycelia. It is often used as an organic amendment to improve soil properties. SMS has certain disease-suppressive functions, particularly showing efficacy against Fusarium-induced crop wilt diseases (Wang et al., 2020).

In earlier work, we demonstrated that amendment with SMS can reduce the incidence of cucumber Fusarium wilt (Qu et al., 2025). This disease is caused by *Fusarium oxysporum* f. sp. *cucumerinum* (FOC), a pathogen that poses a major threat to cucumber cultivation. Its severity is largely due to FOC’s strong infectivity, long-term persistence in soil, and production of FA as a key virulence factor (Liu et al., 2020). SMS does not exhibit direct antifungal activity against FOC but instead mitigates disease development by modulating the cucumber rhizosphere microbiome and enriching beneficial bacteria, such as *Bacillus* (Qu et al., 2025). Although incapable of directly inhibiting the pathogen, SMS reduces the FA concentration in FOC culture media, potentially through adsorption. The toxin-adsorption potential of SMS has been validated in the fields of food and environmental safety. SMS can effectively adsorb various mycotoxins, including aflatoxin B₁ and fumonisins (Kulshreshtha, 2018).

Whether SMS reduces the FA concentration in soil, and if so, how, and whether this is directly associated with the changes in the rhizosphere microbial community and disease suppression, remains unclear. Therefore, in the present study, we investigated whether FA is adsorbed by SMS used as a soil amendment. This included the physicochemical properties of SMS underlying this effect, and the consequences of this adsorption for the rhizosphere microbial community and disease suppression.

## 2. Materials and methods

### 2.1 SMS and microbial Strains

SMS derived from shiitake (*Lentinula edodes*) cultivation was collected from a mushroom production facility in Fuping County, Hebei Province, China. The substrate was air dried for 3 months, ground to a particle size of 1 mm, and sterilised at 121 ℃ for 30 min. To remove water-soluble components, the substrate was suspended in sterile deionised water and shaken at 180 rpm for 1 h. The suspension was centrifuged at 10,000 rpm, and the pellet was resuspended in sterile water. This washing process was repeated five times. Soluble components in the final extract were quantified (Table 1). The treated SMS was oven dried at 80 ℃ for 24 h and ground to a fine powder with a particle size of 100 μm. All the subsequent experiments used SMS after this water-extraction pretreatment.

**Table 1.** Soluble component content of SMS before and after water extraction

| Parameter | Before | After |
| --- | --- | --- |
|  | extraction | extraction |
| pH | 5.05 | 7.05 |
| Electronic conductivity (ds/m) | 4.25 | 0.28 |
| Dissolved total nitrogen, DTN (g/L) | 16.34 | 1.98 |
| Dissolved organic carbon, DOC (g/L) | 10.32 | 1.54 |
| Dissolved phosphorus, DTP (g/L) | 7.19 | 0.84 |
| Dissolved potassium (g/L) | 15.24 | 2.47 |

Bacterial strains used in this study were stored in Luria–Bertani (LB) broth containing 30% (v/v) glycerol at −80 ℃. Before use, the strains were revived by streaking single colonies on LB agar plates and incubating at 37 ℃. The pathogen FOC was also isolated and preserved in our lab. The FOC strain was maintained on potato dextrose agar (PDA) slants at 4 ℃. The mycelia were transferred to PDA plates and incubated at 25 ℃ for 5 d. Agar plugs were then inoculated into potato dextrose broth (PDB) and shaken at 180 rpm, 25 ℃ for 5 d to produce conidia. The resulting culture was filtered to remove mycelia, and the spore suspension was centrifuged at 8000 rpm for 10 min at 4 ℃. The pellet was resuspended in sterile deionised water and adjusted to 10⁷ spores/mL.

### 2.2 Quantification of fusaric acid using high-performance liquid chromatography (HPLC)

FA concentrations in liquid media or soil extracts were quantified using HPLC. The samples were filtered through a 0.2 μm membrane to remove microbial cells and debris. A 10 mL aliquot of the filtrate was mixed with an equal volume of ethyl acetate and shaken for 30 min. The organic phase was separated using a Büchner funnel, and the aqueous phase was discarded. The organic extract was concentrated via rotary evaporation, and the residue was redissolved in 10 mL of methanol as the test solution.

HPLC was performed using a C18 column (250 mm × 4.6 mm, 5 μm) with a mobile phase of methanol:water (60:40, v/v) at a flow rate of 1.0 mL/min and a detection wavelength of 280 nm. A standard curve was constructed using FA standards dissolved in methanol and diluted to 0.1, 1, 10, 50, and 100 μg/mL. The peak areas were used to calculate the FA concentrations in the samples.

### 2.3 Effect of SMS on FA production by FOC

A co-culture experiment was conducted to assess whether SMS affects FA production by FOC. FOC spores (1% v/v, 10⁷ spores/mL) were inoculated into PDB medium containing 1% (w/w) SMS and incubated at 25 ℃, 180 rpm for 7 d. In parallel, an adsorption-only group was prepared by adding 2% SMS (w/w) to SMS-free culture medium after FOC had already grown and incubated for an additional 12 h. The control group (CK) contained no SMS. FA concentration in the culture supernatant was measured using HPLC.

RT-qPCR was used to evaluate the expression of the FA biosynthetic gene *FUB1* in FOC. FOC was cultured on PDA plates containing 1% SMS at 25 ℃ for 3 d. Mycelia were harvested, flash-frozen in liquid nitrogen, and ground into powder. Total RNA was extracted using the RNAprep Pure Cell/Bacteria Kit (TIANGEN, Beijing, China) according to the manufacturer’s instructions. cDNA was synthesized using the FastKing RT Kit (TIANGEN, Beijing, China) following the manufacturer ’s protocol. Quantitative PCR was performed using Talent qPCR PreMix on a 7500 Fast Real-Time PCR System.

*FUB1* expression was quantified using the 2^−ΔΔCT^ method with *actin* as the reference gene. Each treatment included three biological and five technical replicates.

### 2.4 In vitro adsorption of FA by SMS

Adsorption isotherm and kinetic experiments were conducted to evaluate the FA adsorption capacity of SMS. FA solutions (1, 10, 20, 50, and 100 μg/L) were mixed with 1% SMS (w/v) and shaken at 180 rpm, 25 ℃ for 24 h. After incubation, SMS was removed by filtration, and the residual FA concentration in the filtrate was measured. Adsorption isotherms were plotted and fitted using Langmuir and Freundlich models to describe monolayer and heterogeneous adsorption behaviors, respectively (Acharya et al., 2023). For the kinetics analysis, 1% SMS was added to a fixed concentration of FA solution (100 μg/L), and samples were obtained at 1, 2, 4, 8, and 12 h. After filtration, the FA concentrations were measured to plot the kinetic curves. Pseudo-first-order, pseudo-second-order, and Elovich models were applied to fit the kinetic data, describing the adsorption rate, mechanism, and surface interaction processes (Al-Odayni et al., 2023).

### 2.5 Functional group characterisation and chemical modification of SMS

To identify changes in SMS surface functional groups before and after FA adsorption, Fourier Transform Infrared Spectroscopy (FTIR) was performed. SMS (1%) was mixed with 100 μg/mL FA solution and shaken for 1 h at 180 rpm. The mixture was centrifuged at 10,000 rpm, and the pellet was freeze-dried under vacuum. The dried powder was mixed with potassium bromide (KBr) at a 1:100 mass ratio, ground, and pressed into transparent pellets. FTIR spectra were acquired using a Bruker Tensor 27 spectrometer (4000–400/cm, resolution 4/cm, 32 scans). Changes in characteristic absorption peaks were analysed to identify functional group alterations.

To block active functional groups, chemical modification was performed based on (Lee et al., 2009). SMS (1 g) was suspended in 100 mL anhydrous dimethyl sulfoxide containing 0.1 M N-acetylimidazole (NAI) and reacted under nitrogen at 40 ℃ for 4 h. The modified SMS was centrifuged and washed three times with anhydrous acetonitrile, then dried under vacuum for 12 h. FTIR was used to confirm the modification. FA adsorption experiments were conducted using modified and unmodified SMS. Samples were incubated with 100 μg/mL FA for 1 h at 180 rpm. After incubation, supernatants were analysed to determine FA concentrations, and the adsorption efficiency was compared to assess the role of functional groups.

### 2.6 Pot experiment

A dual-compartment pot system (Fig. 3A) was designed to evaluate the effect of SMS-mediated FA adsorption on cucumber Fusarium wilt suppression. Nylon mesh bags (48 μm pore size) were used to separate the inner and outer compartments of each pot. The outer compartment was filled with natural soil mixed with 1% (w/w) SMS. Meanwhile, the inner compartment contained untreated soil in which one cucumber s (*Cucumis sativus* L. cv. Jinyan No. 4) eedling was planted per pot.

**Figure 1.**
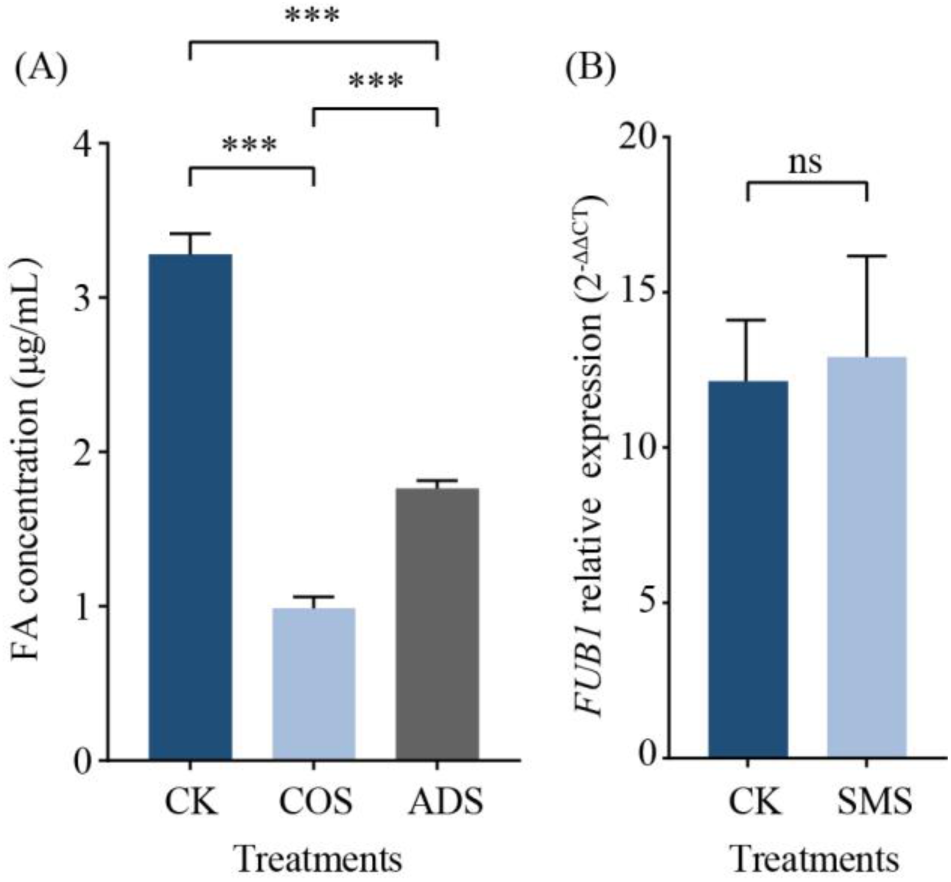
Effects of SMS on FA production and FUB1 gene expression in FOC. (A) FA concentrations in culture supernatants under CK (control), COS (SMS co-incubated with FOC), and ADS treatments with SMS added after FOC culture. (B) Relative expression of the FA biosynthetic gene FUB1 under CK and SMS treatments.

When the seedlings reached the two-leaf stage, 100 mL of FA solution at low (20 μg/mL), medium (40 μg/mL), and high (80 μg/mL) concentrations was applied to the outer compartment soil. A no-SMS control was included. One hour later, 1 g of soil from the inner compartment was collected for FA concentration analysis. After flushing with sterile deionised water to reduce residual FA to <1 μg/mL, 50 mL of FOC spore suspension (10⁷ spores/mL) was introduced into the inner compartment. Disease assays included three treatments: (1) CK: untreated natural soil; (2) SMS: outer compartment with 1% SMS-amended soil; and (3) Sterilised: outer compartment with sterilised SMS-amended soil. Disease severity was scored after 7 d using a five-grade scale (Zhou et al., 2017), and the disease index was calculated. Each treatment had three replicates with 18 plants per replicate.

### 2.7 Microbial diversity analysis

After the pot experiment, the cucumber seedlings were gently uprooted, and loosely attached soil was removed. The rhizosphere soil was collected using sterile brushes, sieved through a 50-mesh screen, and stored at −80 ℃. DNA was extracted from 0.5 g of rhizosphere soil using the FastDNA™ SPIN Kit (MP Biomedicals, USA) according to the manufacturer’s instructions. DNA concentration and purity were assessed with a NanoDrop 2000 spectrophotometer (Thermo Fisher Scientific, USA). The V3–V4 region of the 16S rRNA gene was amplified using primers 338F (5′-ACTCCTACGGGAGGCAGCAG-3′) and 806R (5′-GGACTACHVGGGTWTCTAAT-3′). PCR amplification, product purification, and library preparation were performed following the method described by (Zhang et al., 2019). Sequencing was performed on the Illumina MiSeq platform (Personal Biotechnology, China), and data analysis was conducted using the Majorbio Cloud platform (Ren et al., 2022).

### 2.8 FA Tolerance of rhizosphere bacteria

To assess the FA tolerance of rhizosphere biocontrol strains, six laboratory-preserved strains were tested: *Bacillus subtilis* NCD-2, *B. velezensis* GY2-11, *B. velezensis* MTC-28, *B. subtilis* MTS-27, *Pseudomonas fluorescens* 2P24, and *Streptomyces griseoviridis* R-24. Each strain was pre-cultured in 5 mL LB broth at 37 ℃, 180 rpm for 12 h. Then, 100 μL of the culture was inoculated into 50 mL of LB medium containing different FA concentrations: negative control (NC) = 0 μg/mL, F-L = 10 μg/mL, F-M = 20 μg/mL, and F-H = 40 μg/mL. Cultures were incubated at 37 ℃, 180 rpm for 36 h. OD600 values were measured at 0, 2, 4, 6, 8, 24, and 36 h to construct growth curves and evaluate the FA tolerance.

## 3. Results

### 3.1 Effect of SMS on FA production by FOC

A flask co-culture experiment was conducted to assess the influence of SMS on FA production by FOC. FA concentrations in the co-culture group (COS, SMS co-incubated with FOC) and the adsorption group (ADS, SMS added after FOC culture) were 1.39 μg/mL and 1.76 μg/mL, respectively. This represented reductions of 57.62% and 46.34% compared to the control group (CK, 3.28 μg/mL). SMS significantly decreased the FA concentration in the culture supernatant, whether co-incubated with FOC or added afterwards. RT-qPCR analysis showed no significant change in the expression level of the FA biosynthetic gene *FUB1* in response to the SMS treatment. The results suggest that the reduction in the FA concentration in the culture supernatant is likely attributable to adsorption of FA by SMS rather than suppression of FA biosynthesis.

### 3.2 Adsorption characteristics of SMS towards FA

To evaluate the adsorption behaviour of SMS toward FA, adsorption isotherms were generated by measuring the equilibrium adsorption capacity (*Qe*) at different initial FA concentrations (*Ce*). The adsorption capacity increased with rising FA concentrations and approached saturation. Model fitting showed that the Langmuir–Freundlich composite model best described the isotherm, with a theoretical maximum adsorption capacity (*Qm*) of 52.72 μg/L. This indicated the involvement of both physical and chemical adsorption mechanisms (Fig. 2A, Table 2). Adsorption kinetics were further analysed using nonlinear fitting. The process was best described by the pseudo-second-order kinetic model (*R²* = 0.78), with an equilibrium adsorption capacity (*Qt*) of 48.92 μg/g (Fig. 2B, Table 2). These results suggest that FA adsorption by SMS involved chemical interaction and physical entrapment. Scanning electron microscopy indicated that SMS possessed a porous surface structure (Fig. 2C), which likely facilitated efficient mass transfer and rapid physical adsorption of FA molecules.

**Figure 2.**
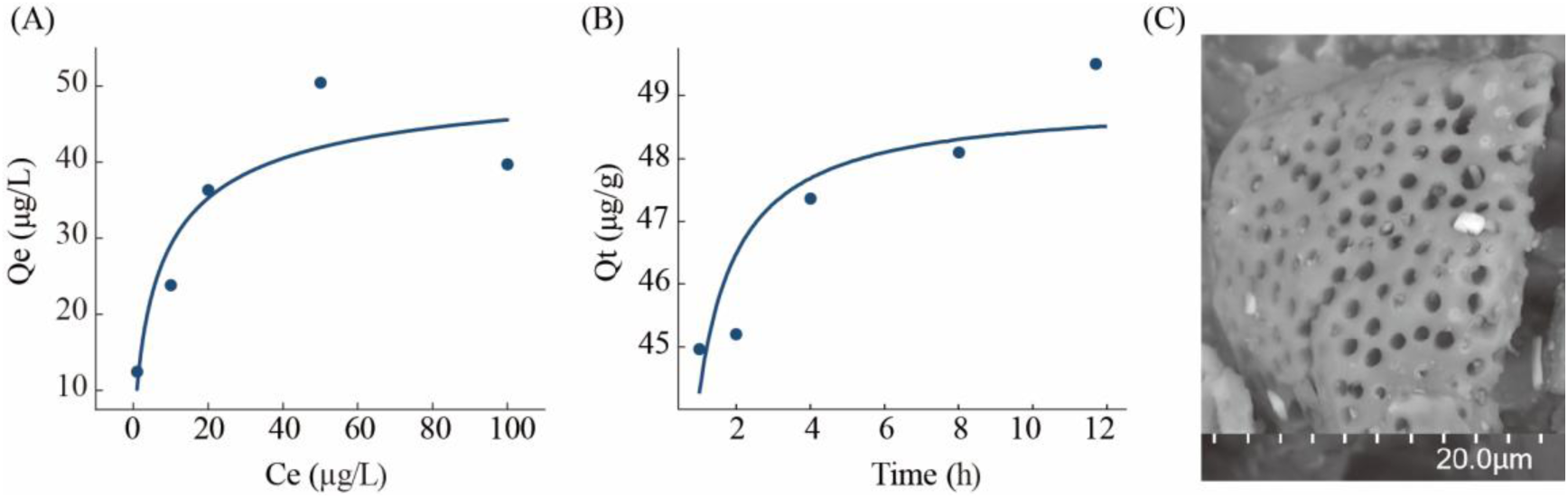
Adsorption of FA by SMS: Langmuir–Freundlich isotherm fitting (A), pseudo-second-order kinetic fitting (B), and SEM observation of SMS surface morphology (C).

**Figure 3.**
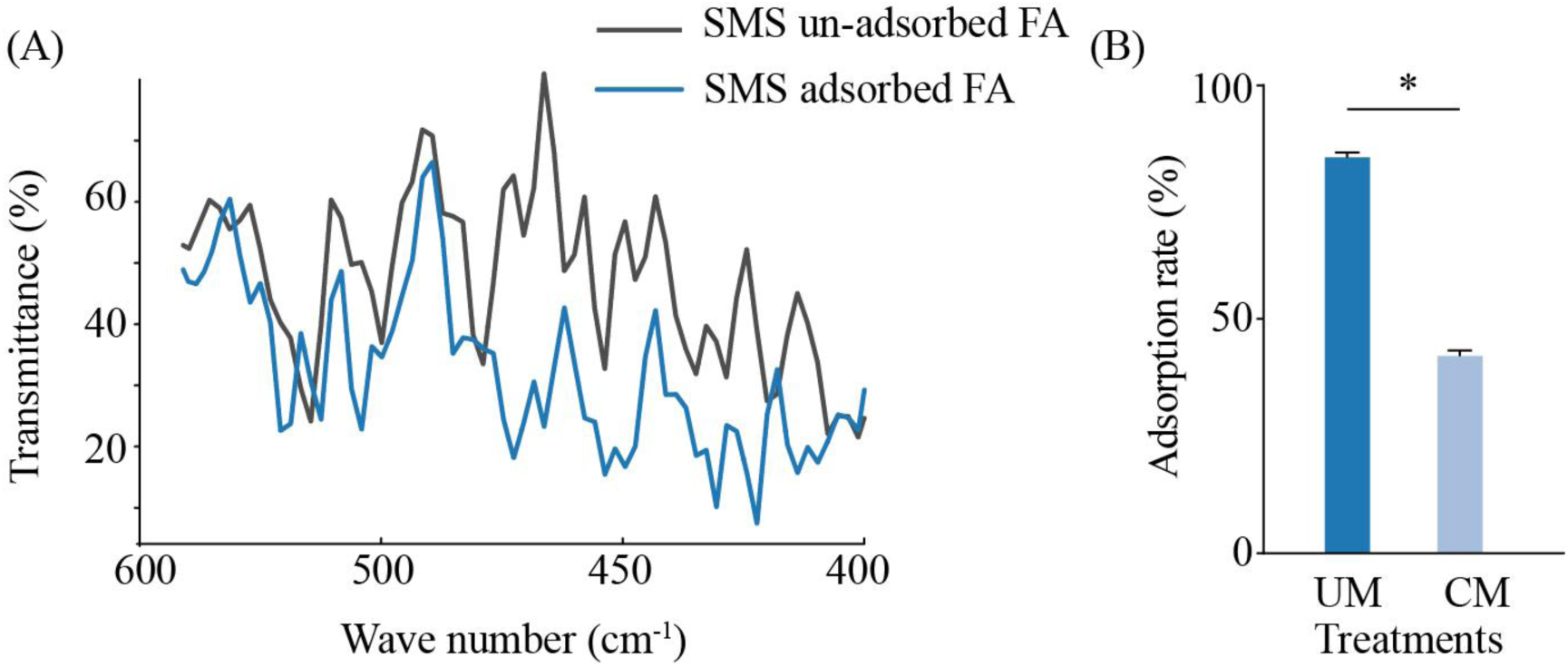
Functional group analysis of SMS in FA adsorption. (A) FTIR spectra of SMS before and after FA adsorption. (B) FA adsorption rates of unmodified (UM) and CM SMS, showing reduced efficiency after acetylation.

**Table 2.**
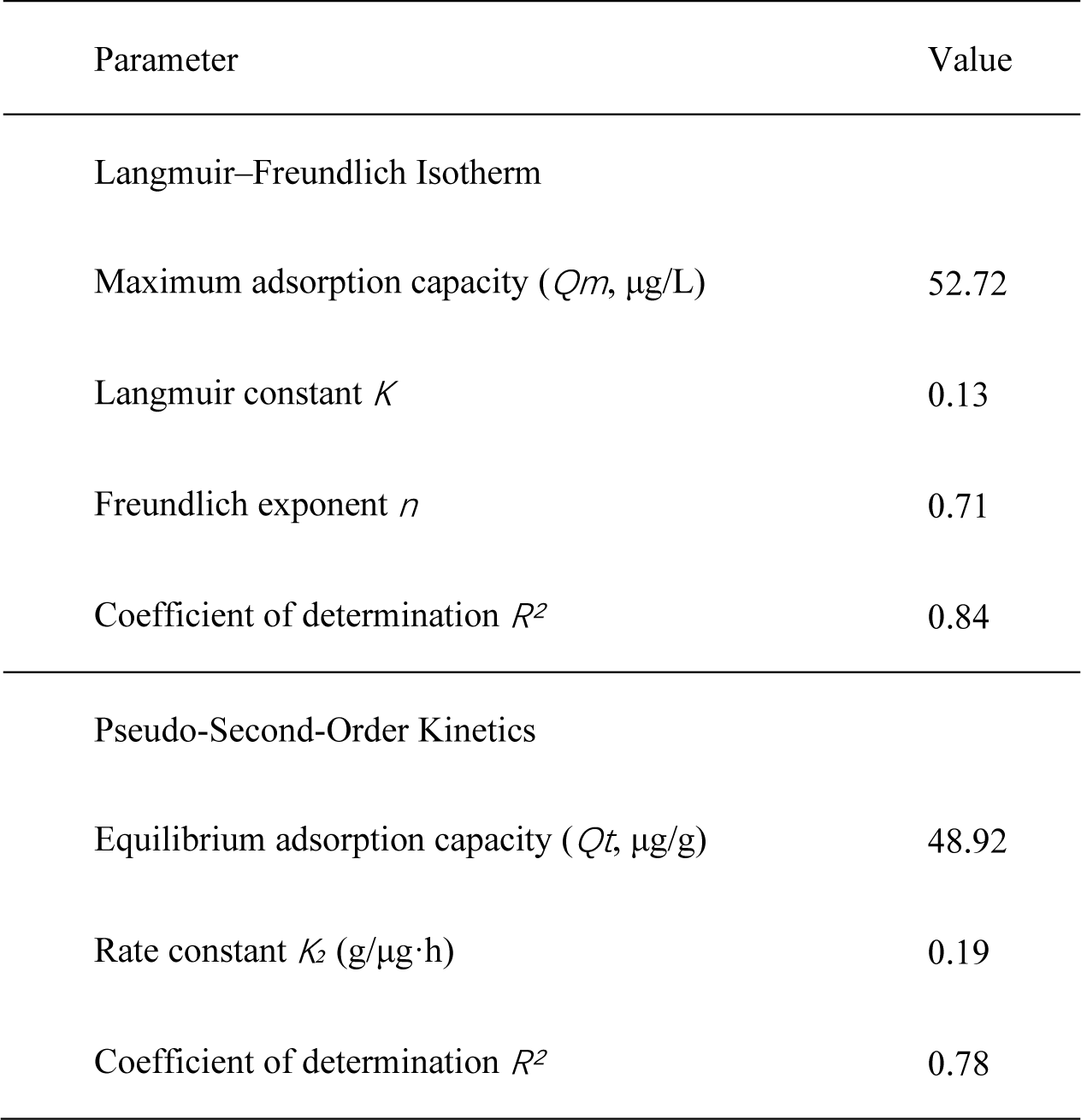
Adsorption parameters of SMS for FA

| Parameter | Value |
| --- | --- |
| Langmuir–Freundlich Isotherm |  |
| Maximum adsorption capacity ($Q_m$, $\mu\text{g/L}$) | 52.72 |
| Langmuir constant $K$ | 0.13 |
| Freundlich exponent $n$ | 0.71 |
| Coefficient of determination $R^2$ | 0.84 |
| Pseudo-Second-Order Kinetics |  |
| Equilibrium adsorption capacity ($Q_t$, $\mu\text{g/g}$) | 48.92 |
| Rate constant $K_2$ ($\text{g}/\mu\text{g}\cdot\text{h}$) | 0.19 |
| Coefficient of determination $R^2$ | 0.78 |

### 3.3 Functional group analysis

To elucidate the chemical interactions involved in FA adsorption, FTIR analysis was conducted to examine changes in SMS surface functional groups before and after adsorption. The absorption peak corresponding to the C=O stretching vibration of secondary amide groups (495.6–401.1/cm) decreased after FA adsorption, suggesting that these groups were involved in chemical binding. This interaction was likely mediated by hydrogen bonding or covalent bonding between the carboxyl or phenolic hydroxyl groups of FA and the secondary amide groups on the SMS surface.

To verify this, secondary amide groups on SMS were inactivated via acetylation using NAI. Chemically modified SMS (CM) showed significantly lower FA adsorption than unmodified SMS (UM) (Fig. 3), with the adsorption efficiency being reduced by 45% (*p* < 0.05), indicating that secondary amide groups played a pivotal role in chemical binding. CM retained some adsorption capacity, indicating that FA adsorption by SMS was also driven by physical mechanisms.

### 3.4 Impact of SMS on FOC pathogenicity

A dual-compartment pot system (Fig. 4A) was used to assess the capacity of SMS to adsorb FA in soil and whether that would mitigate its pathogenic effects. After FA solutions were applied to the outer compartment soil, the inner compartment was analysed for FA. In the control group (CK) without SMS, the FA concentration in the inner compartment reached 15.75, 31.50, and 62.29 μg/g under low (LF, 20 μg/mL), medium (MF, 40 μg/mL), and high FA (HF, 80 μg/mL) treatments, respectively. In contrast, the corresponding FA concentrations in the SMS treatment group, with 1% SMS added to the outer compartment, were significantly reduced to 3.25, 8.31, and 15.49 μg/g. This confirmed that SMS could adsorb FA in soil (Fig. 4B).

**Figure 4.**
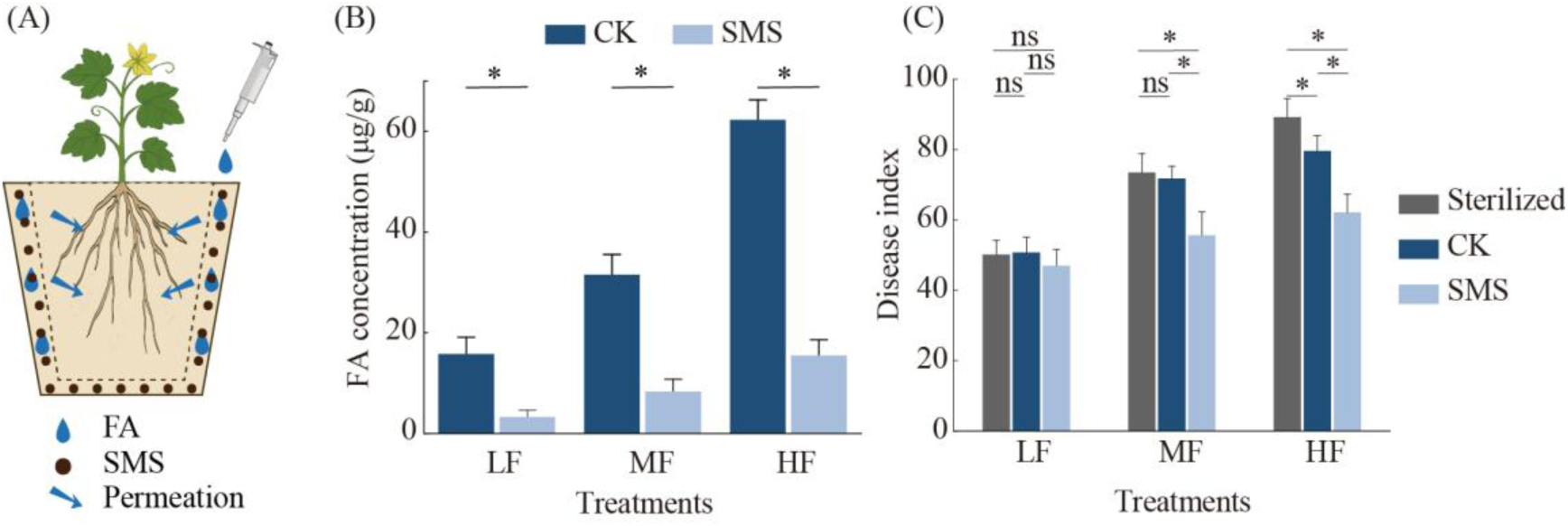
Effects of SMS on FA adsorption in soil and suppression of FOC pathogenicity in a dual-compartment pot system. (A) Experimental setup of the dual-compartment pot system. (B) FA concentrations in the inner soil compartment under low (LF), medium (MF), and high (HF) FA treatments with or without SMS. (C) Disease indices of cucumber seedlings under CK, SMS, and sterilised SMS treatments.

The same system was used to evaluate the effect of SMS and FA on disease severity 7 d after inoculation of FOC. In the CK group, disease indices for MF and HF treatments increased by 42% and 57% compared to LF, respectively. This suggested that higher FA concentrations enhanced FOC virulence. However, in the SMS-treated group, these increases were partially mitigated to 19% and 32%, respectively. Relative to CK, disease indices were reduced by 23% and 25% under the MF and HF treatments, respectively. This indicated that SMS adsorption alleviated FA-induced disease aggravation. In the sterilised SMS treatment (Sterilised), where soil microbiota was inactivated, disease indices significantly increased compared to the non-sterilised SMS treatment by 31.94% and 43.61% under the MF and HF treatments, respectively. This suggests that SMS-mediated FA adsorption has a more pronounced protective effect in biologically active soils (Fig. 4C).

### 3.5 Rhizosphere bacterial community analysis

To further investigate the role of soil microbes in SMS-mediated disease suppression through FA adsorption, the rhizosphere bacterial community was analysed via 16S rRNA gene amplicon sequencing.

A total of 46 bacterial phyla were annotated in the cucumber rhizosphere samples. The dominant phyla were *Proteobacteria* (48.00%), followed by *Bacteroidetes* (14.70%), *Actinobacteria* (13.70%),

*Acidobacteria* (8.47%), *Verrucomicrobia* (4.00%), *Chloroflexi* (3.20%), *Gemmatimonadetes* (3.11%), and *Firmicutes* (1.22%). The taxa with relative abundances below 1% were grouped as ‘others’.

Compared with the negative control (NC), medium (F-M) and high (F-H) FA treatments significantly reduced bacterial diversity and richness. This suggests that FA disrupted the microbial community structure. In contrast, the SMS-adsorbed medium (S-F-M) and high (S-F-H) FA treatments showed no significant difference in diversity or richness compared to NC. This suggests that SMS effectively mitigated the toxic effects of FA. At low FA concentrations, neither FA nor SMS treatment significantly affected the microbial diversity. β-diversity and heatmap clustering analyses further supported these findings. F-M and F-H formed distinct clusters, indicating community shifts. Meanwhile, S-F-M, S-F-H, S-F-L, and F-L clustered closely with NC, suggesting that SMS reduced FA-induced community disruption (Fig. 5).

**Figure 5.**
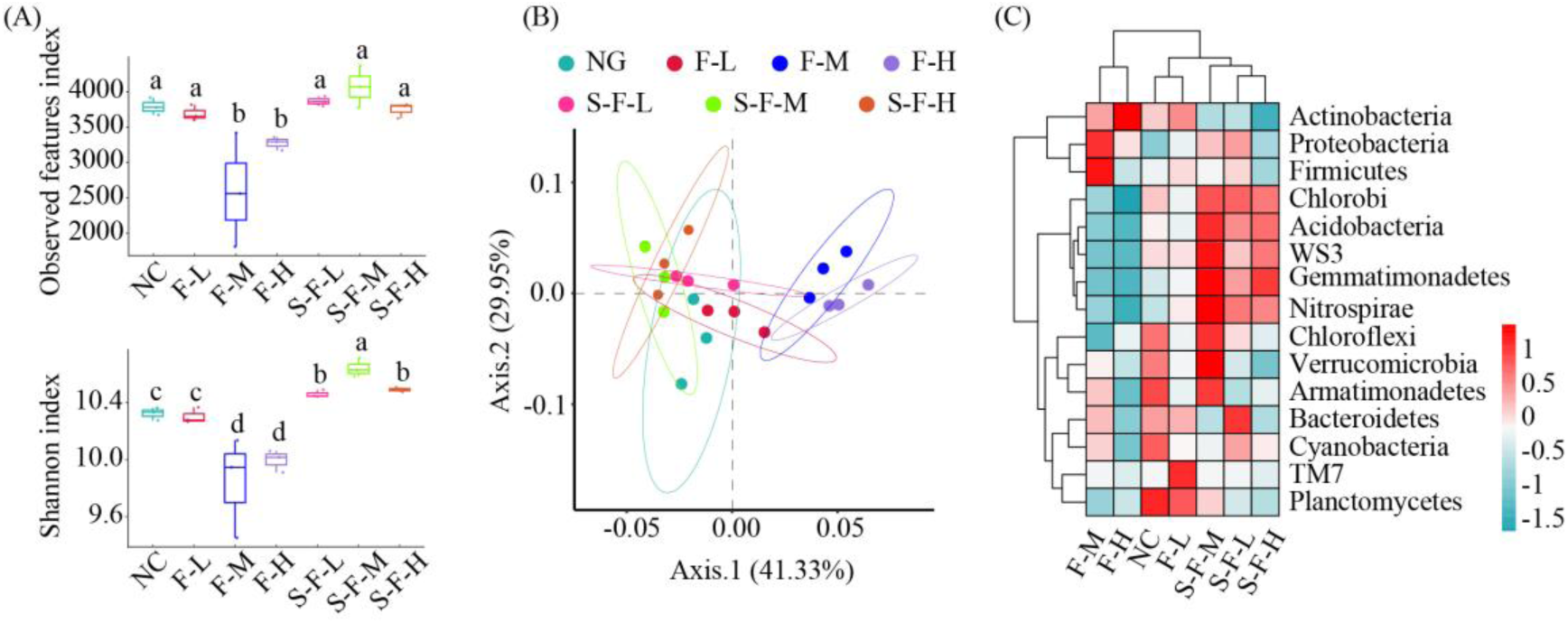
Rhizosphere bacterial community analysis under FA and SMS treatments. (A) Observed features index and Shannon index. (B) Principal coordinate analysis (PCoA) of bacterial β-diversity. (C) Heatmap of dominant bacterial phyla.

To identify taxonomic responses at the genus level, the relative abundance of bacterial genera was compared among the NC, F-H, and S-F-H groups. A total of 239 genera were annotated, with the top five being *Streptomyces* (4.17%), *Rhodanobacter* (3.34%), *Kaistobacter* (3.06%), *Bacillus* (2.78%), and *Flavisolibacter* (2.15%). Under the F-H treatment, the relative abundance of 19 genera declined significantly compared to NC, indicating high FA sensitivity. Of these, seven genera exhibited significantly higher abundances in S-F-H than in F-H, suggesting that SMS adsorption alleviated FA toxicity. For example, *Bacillus* declined from 3.18% (NC) to 2.19% (F-H) but was maintained at 2.96% in the S-F-H group, showing no significant difference from NC. In contrast, *Streptomyces* and *Pseudomonas* were unaffected by FA stress, indicating greater intrinsic tolerance (Fig. 6).

**Figure 6.**
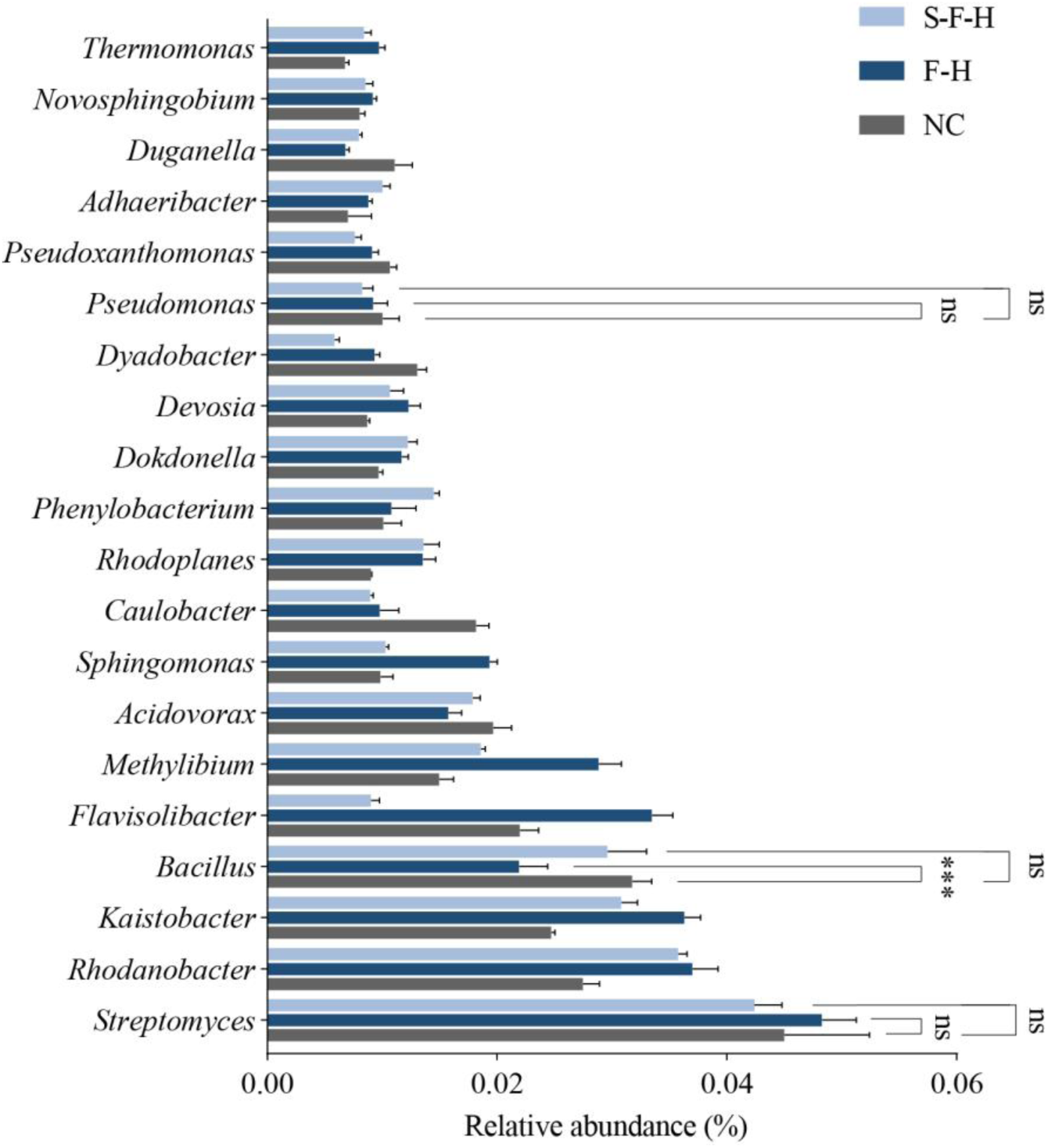
Relative abundances of dominant bacterial genera in cucumber rhizosphere under NC, F-H, and S-F-H treatments.

### 3.6 Tolerance of rhizosphere bacteria to FA

To further underpin the existence of differences in FA tolerance between bacterial species, six representative rhizosphere strains were tested under varying FA concentrations (Fig. 7). *Bacillus subtilis* NCD-2, *B. velezensis* GY2-11, *B. velezensis* MTC-28, and *B. subtilis* MTS-27 showed no significant growth inhibition under F-L (10 μg/mL) and F-M (20 μg/mL) conditions. However, at F-H (40 μg/mL), NCD-2 exhibited substantial growth suppression, with OD600 values of 0.11, 0.11, 0.23, and 0.25 at 8, 12, 24, and 36 h, respectively, representing reductions of 38.89%, 75.00%, 79.46%, and 83.22%, respectively, compared to the NC group (OD600: 0.18, 0.44, 1.12, and 1.49). Similar trends were observed for the other three *Bacillus* strains (GY2-11, MTC-28, and MTS-27). In contrast, *Pseudomonas fluorescens* 2P24 and *Streptomyces griseoviridis* R-24 showed no significant differences in growth between the F-H and NC treatments. These findings demonstrate that *Bacillus* spp. was sensitive to FA, while *Pseudomonas* and *Streptomyces* exhibited tolerance.

**Figure 7.**
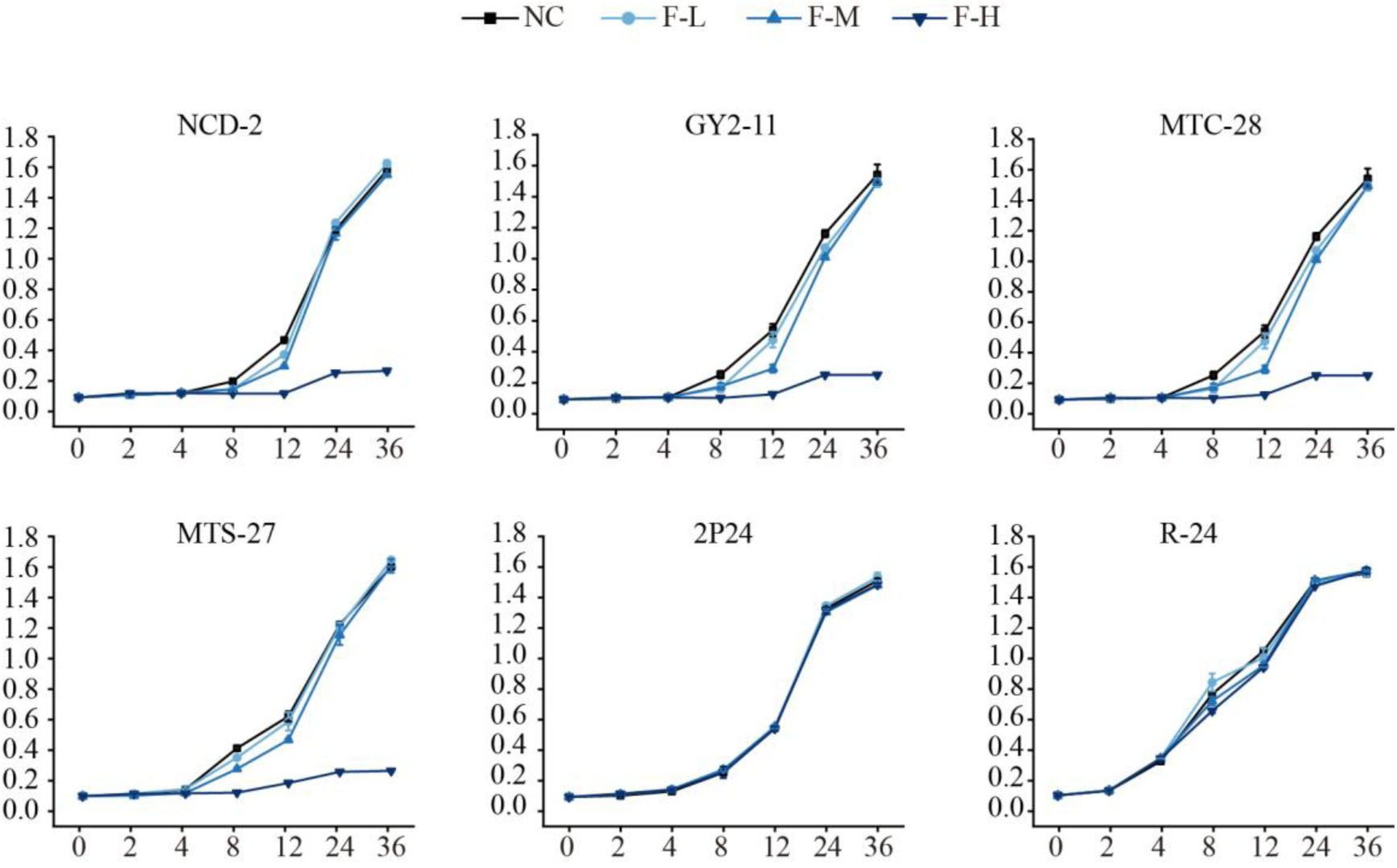
Growth curves of six bacterial strains under different FA concentrations (F-L: 10 μg/mL; F-M: 20 μg/mL; F-H: 40 μg/mL).

## 4. Discussion

In the present study, we demonstrated that SMS derived from *Lentinula edodes* substantially reduced the bioavailable concentration of FA in soil through adsorption, alleviating its toxic effects on sensitive beneficial bacterial taxa. This, in turn, stabilised the rhizosphere microecosystem and indirectly enhanced plant disease resistance. These findings indicate a novel disease suppression mechanism of SMS, characterised by the pathway of adsorption-based detoxification → microbial protection → disease suppression. This provides a theoretical foundation and practical potential for the ecological regulation of mycotoxins and sustainable plant disease management.

Toxic metabolites secreted by pathogenic fungi are a key strategy for competing for ecological niches in the rhizosphere and suppressing the defensive capacity of beneficial microorganisms (Lu et al., 2025). As a representative mycotoxin produced by *F. oxysporum*, FA exhibits broad-spectrum toxicity. It directly harms plants and disrupts rhizosphere microbial communities, enhancing the ecological competitiveness of the pathogen (Jin et al., 2024). FA has pronounced toxicity on a range of soil microorganisms and can impair their ecological fitness in the rhizosphere through multiple mechanisms (Bacon et al., 2004). FA significantly inhibits the growth and root colonisation of PGPR, such as *Bacillus* spp., reducing their competitiveness for ecological niches and creating spatial advantages for *Fusarium* infection (Prigigallo et al., 2022). Meanwhile, FA acts as an interspecies signalling molecule, interfering with the metabolic functions of PGPR. (Quecine et al., 2016) found that FA inhibits the synthesis of the antifungal metabolite DAPG in *Pseudomonas protegens* Pf-5, weakening its biocontrol potential. FA can disrupt the enrichment and assembly of specific beneficial taxa, such as disease-suppressive microbes, at the plant root interface (Jin et al., 2024). Our study confirmed the toxicity of FA at the community level, with approximately 60 μg FA/g soil leading to a substantial reduction in the diversity and richness of the cucumber rhizosphere bacterial community. This suggests that FA’s ecological interference extends beyond individual strains and compromises the structure and stability of the entire rhizosphere microbiome. Mitigating FA’s microbial toxicity could contribute to maintaining rhizosphere ecological balance and achieving sustainable disease control (Xie et al., 2019).

Adsorption-based techniques have been widely applied for mycotoxin mitigation, especially in the fields of food safety and environmental remediation. Here, they effectively reduce toxin bioavailability and associated toxicity (Ma et al., 2023; Yang et al., 2023). This strategy has been proposed for the ecological regulation of mycotoxins in soil-borne pathosystems. (Jaiswal et al., 2018) reported that biochar can alleviate tomato Fusarium wilt by adsorbing fungal cell wall-degrading enzymes and toxins and proposed an adsorption–inactivation mechanism for disease suppression. SMS adsorbs a range of environmental contaminants, including heavy metals, pesticide residues, and mycotoxins (Alvarez-Martín et al., 2016; Sahithya et al., 2022; Rasheed et al., 2024).

To investigate whether adsorption also plays a role in the FOC suppressive effect of SMS amendment, we used a pot experiment to physically isolate SMS from the plant. Soluble nutrients in SMS were removed through multiple rounds of aqueous extraction to exclude nutritional confounding effects. Under these controlled conditions, the rhizosphere bacterial diversity and community structure of cucumber remained stable in the presence of FA when SMS was applied and the FA was absorbed, in contrast to the FA-only treatment. The adsorption function of SMS likely creates a functional buffer zone between pathogenic and beneficial microbes, mitigating the negative effect of FA on PGPR. This protects the rhizosphere microbiome integrity, enhancing system stability.

There were differences in FA tolerance among rhizosphere bacterial taxa. FA significantly reduced the relative abundance of *Bacillus* spp., while having negligible effects on *Pseudomonas* and *Streptomyces* spp. In treatments where SMS adsorbed FA, the abundance of *Bacillus* spp. was restored to control levels, suggesting a clear selective protection effect. *Pseudomonas* and *Streptomyces* strains are typically equipped with efficient detoxification mechanisms, such as efflux pumps and redox enzymes that reduce intracellular toxin accumulation (Crutcher et al., 2017). Meanwhile, *Bacillus* spp. lack such systems and are prone to growth inhibition and membrane damage, including at low FA concentrations (Bacon et al., 2006). SMS adsorption likely preferentially alleviates toxicity for sensitive taxa, preserving the ecological roles of keystone functional groups. Combined with our previous findings that SMS enriches *Bacillus* populations and suppresses Fusarium wilt (Qu et al., 2025), we propose that FA adsorption by SMS protects *Bacillus* spp. through FA adsorption. This represents a key mechanism in the disease-suppressive effect of SMS. However, such protection primarily helps maintain the ecological niche of sensitive bacteria. Further enrichment of these beneficial taxa likely depends on additional factors, such as the plant root exudate and environmental conditions, which warrant further investigation (Bouwmeester et al., 2025).

The adsorption of FA by SMS is driven by both physical and chemical interactions. The porous surface structure of SMS provides abundant sites for physical entrapment of FA molecules, forming the basis for rapid toxin capture (Kokuloku et al., 2023). Enhancing pore volume and surface area substantially improves the adsorption capacity of porous materials, such as biochar, for organic pollutants (Ullah and Rahman, 2024). In our study, SMS also had abundant porosity. FTIR analysis indicated the presence of phenolic hydroxyl, carboxyl, and secondary amide groups on the SMS surface, highlighting the potential for chemical bonding with FA. Functional group-blocking experiments suggested that these groups acted co-operatively in the chemical adsorption process. Kinetic analysis showed that the adsorption followed a pseudo-second-order model, indicating that the rate-limiting step involved chemical bonding. Isotherm fitting results aligned better with the Langmuir–Freundlich composite model. This suggests that the SMS surface is heterogeneous and that adsorption is driven by a combination of physical capture and chemical interaction. These results provide mechanistic insights into the adsorption of FA by SMS and offer theoretical guidance for the optimisation of organic adsorbents to be used in agriculture for sustainable plant disease management.

FA was uniformly applied in solution, and adsorption was regulated using physical isolation to focus on the adsorption mechanism. However, in field environments, FA is intermittently secreted by pathogens and exhibits uneven spatial distribution, with its behaviour influenced by multiple ecological factors. Whether SMS can effectively adsorb FA under field conditions and mitigate its ecological toxicity remains to be demonstrated using field experiments. Future research should focus on the in-situ release dynamics of FA and assess the adsorption capacity and disease control efficacy of SMS across different soil types and cropping systems to enhance its generalizability and reliability in agricultural applications.

In this study, we examined the mechanism by which *Lentinula edodes* SMS adsorbs FA, alleviating its ecological toxicity (Fig. 8). This FA adsorption results in the stabilisation of the rhizosphere microbial community, particularly by protecting sensitive *Bacillus* PGPRs and suppressing soil-borne disease. The SMS porous surface structure provides physical adsorption sites for FA. Meanwhile, functional groups, such as secondary amides, contribute to chemical bonding and facilitate efficient FA immobilisation. This adsorption process substantially reduces the bioavailable concentration of FA in the rhizosphere and alleviates its toxicity toward sensitive beneficial taxa, such as *Bacillus*. It also maintains the stability of the rhizosphere microbiome while weakening the ecological advantage of the pathogen. Our findings have established and validated a mechanistic pathway for the adsorption-based detoxification of pathogen-produced toxins by organic amendments that result in microbiome protection and disease suppression. This has shown a mechanistic cascade through which SMS mediates ecological mitigation of plant diseases in the rhizosphere (Fig. 8). This mechanism advances our understanding of the ecological roles of mycotoxins and provides theoretical and technical support for the valorisation of SMS and the development of sustainable strategies for managing soil-borne plant diseases.

**Figure 8.**
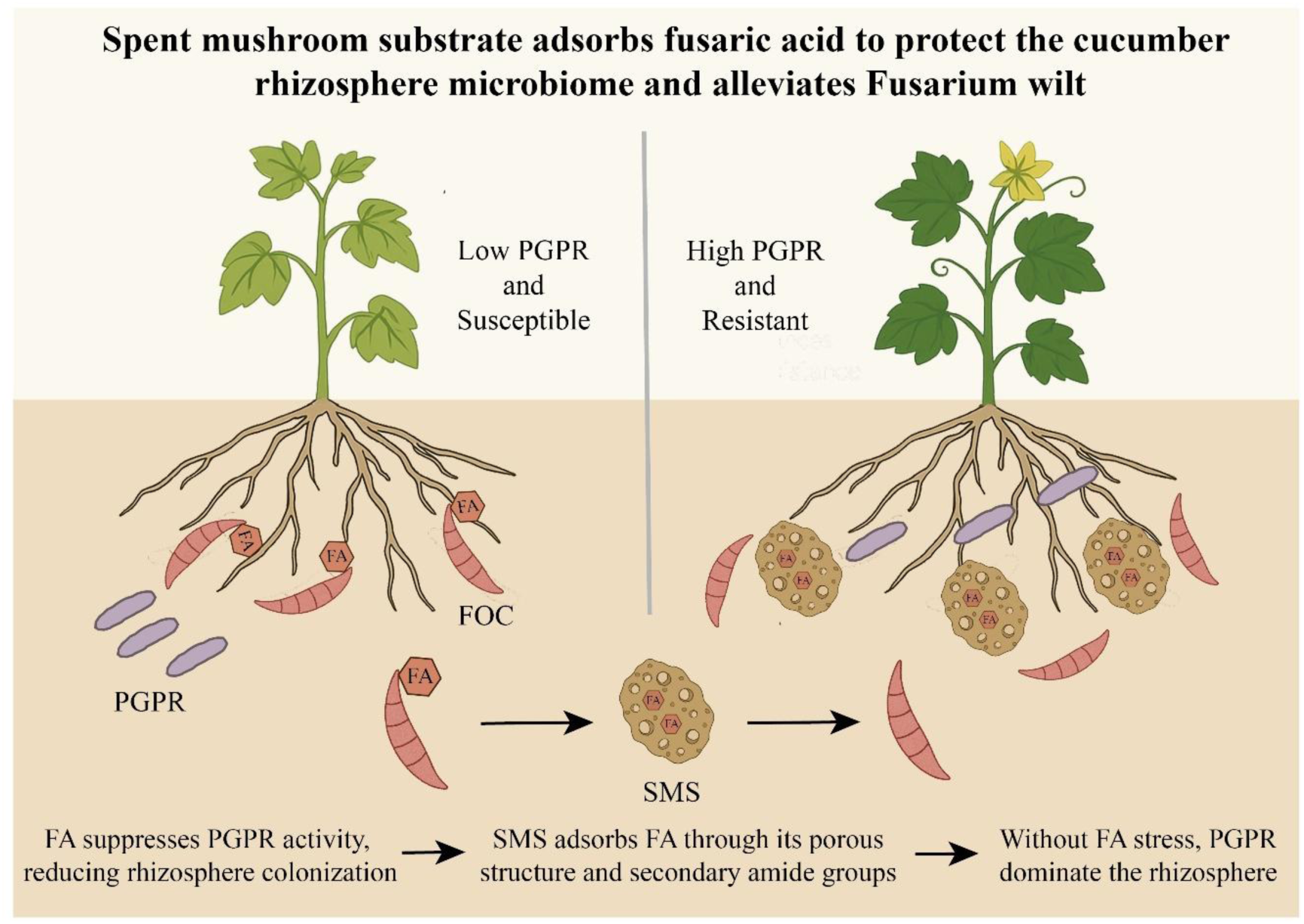
Proposed mechanism of SMS-mediated alleviation of cucumber Fusarium wilt. FA: fusaric acid; PGPR: plant growth-promoting rhizobacteria; FOC: Fusarium oxysporum f. sp. Cucumerinum; SMS: spent mushroom substrate.

## References

1. Acharya, A., Jeppu, G., Raju Girish, C., Prabhu, B., 2023. Development of a multicomponent adsorption isotherm equation and its validation by modeling. Langmuir 39, 17862–17878.

2. Al-Odayni, A.-B., Alsubaie, F.S., Abdu, N.A., Al-Kahtani, H.M., Saeed, W.S., 2023. Adsorption kinetics of methyl orange from model polluted water onto N-doped activated carbons prepared from N-containing polymers. Polymers 15, 1983.

3. Alvarez-Martín, A., Sanchez-Martín, M., Pose-Juan, E., Rodríguez-Cruz, M., 2016. Effect of different rates of spent mushroom substrate on the dissipation and bioavailability of cymoxanil and tebuconazole in an agricultural soil. Science of the Total Environment 550, 495–503.

4. Bacon, C., Hinton, D., Hinton, A., 2006. Growth-inhibiting effects of concentrations of fusaric acid on the growth of *Bacillus mojavensis* and other biocontrol Bacillus species. Journal of Applied Microbiology 100, 185–194.

5. Bacon, C., Hinton, D., Porter, J., Glenn, A., Kuldau, G., 2004. Fusaric acid, a *Fusarium verticillioides* metabolite, antagonistic to the endophytic biocontrol bacterium Bacillus mojavensis. Canadian Journal of Botany-Revue Canadienne De Botanique 82, 878–885.

6. Bouwmeester, H., Dong, L., Wippel, K., Hofland, T., Smilde, A., 2025. The chemical interaction between plants and the rhizosphere microbiome. Trends in Plant Science 30, 1002–1019.

7. Brown, M., Jayaweera, D., Hunt, A., Woodhall, J., Ray, R., 2021. Yield Losses and Control by Sedaxane and Fludioxonil of Soilborne Rhizoctonia, Microdochium, and Fusarium Species in Winter Wheat. Plant Disease 105, 2521–2530.

8. Crutcher, F., Puckhaber, L., Stipanovic, R., Bell, A., Nichols, R., Lawrence, K. et al., 2017. Microbial Resistance Mechanisms to the Antibiotic and Phytotoxin Fusaric Acid. Journal of Chemical Ecology 43, 996–1006.

9. Fu, C., Zhang, H., Xia, M., Lei, W., Wang, F., 2020. The single/co-adsorption characteristics and microscopic adsorption mechanism of biochar-montmorillonite composite adsorbent for pharmaceutical emerging organic contaminant atenolol and lead ions. Ecotoxicology and Environmental Safety 187, 109763.

10. Jaiswal, A., Frenkel, O., Tsechansky, L., Elad, Y., Graber, E., 2018. Immobilization and deactivation of pathogenic enzymes and toxic metabolites by biochar: A possible mechanism involved in soilborne disease suppression. Soil Biology & Biochemistry 121, 59–66.

11. Jin, X., Jia, H., Ran, L., Wu, F., Liu, J., Schlaeppi, K. et al., 2024. Fusaric acid mediates the assembly of disease-suppressive rhizosphere microbiota via induced shifts in plant root exudates. Nature Communications 15, 5125.

12. Kokuloku, L.J., Miensah, E., Gu, A., Chen, K., Wang, P., Gong, C. et al., 2023. Efficient and comparative adsorption of trinitrotoluene on MOF MIL-100 (Fe)-derived porous carbon/Fe composite adsorbents with rod-like morphology: Behavior, mechanism, and new perspectives. Colloids and Surfaces a-Physicochemical and Engineering Aspects 663, 131064.

13. Kulshreshtha, S., 2018. Mushroom biomass and spent mushroom substrate as adsorbent to remove pollutants, Green adsorbents for pollutant removal: innovative materials. Springer, pp. 281–325.

14. Lee, J., Farha, O.K., Roberts, J., Scheidt, K.A., Nguyen, S.T., Hupp, J.T., 2009. Metal–organic framework materials as catalysts. Chemical Society Reviews 38, 1450–1459.

15. Li, Y., Tian, G., Dong, G., Bai, S., Han, X., Liang, J. et al., 2018. Research progress on the raw and modified montmorillonites as adsorbents for mycotoxins: A review. Applied Clay Science 163, 299–311.

16. Liu, S., Li, J., Zhang, Y., Liu, N., Viljoen, A., Mostert, D. et al., 2020. Fusaric acid instigates the invasion of banana by *Fusarium oxysporum* f. sp. *cubense* TR4. New Phytologist 225, 913–929.

17. Lu, H., Guo, S., Yang, Y., Zhao, Z., Xie, Q., Wu, Q. et al., 2025. Bikaverin as a molecular weapon: enhancing F*usarium oxysporum* pathogenicity in bananas via rhizosphere microbiome manipulation. Microbiome 13, 107.

18. Ma, F., Guo, Q., Zhang, Z., Ding, X., Zhang, L., Li, P. et al., 2023. Simultaneous removal of aflatoxin B1 and zearalenone in vegetable oils by hierarchical fungal mycelia@graphene oxide@Fe3O4 adsorbent. Food Chemistry 428, 136779.

19. Noman, M., Ahmed, T., Islam, M., Wang, J., Cai, Y., Liang, S. et al., 2025. Bacterial extracellular biomolecules-derived multimodal manganese nanoparticles control watermelon Fusarium wilt by dysregulating fusaric acid biosynthesis pathway and precise tuning of rhizosphere metabolome. Journal of Nanobiotechnology 23, 452.

20. Notz, R., Maurhofer, M., Dubach, H., Haas, D., Défago, G., 2002. Fusaric acid-producing strains of *Fusarium oxysporum* alter 2,4-diacetylphloroglucinol biosynthetic gene expression in *Pseudomonas fluorescens* CHA0 in vitro and in the rhizosphere of wheat. Applied and Environmental Microbiology 68, 2229–2235.

21. Prigigallo, M., Cabanas, C., Mercado-Blanco, J., Bubici, G., 2022. Designing a synthetic microbial community devoted to biological control: The case study of Fusarium wilt of banana. Frontiers in Microbiology 13, 967885.

22. Qu, Y., Liu, T., Dong, L., Dong, L., Su, Z., Bouwmeester, H. et al., 2025. Spent mushroom substrate amendment induces suppressiveness against cucumber Fusarium wilt through changes in the rhizosphere microbiome. Plant And Soil 514, 3043–3058.

23. Quecine, M., Kidarsa, T., Goebel, N., Shaffer, B., Henkels, M., Zabriskie, T. et al., 2016. An Interspecies Signaling System Mediated by Fusaric Acid Has Parallel Effects on Antifungal Metabolite Production by *Pseudomonas protegens* Strain Pf-5 and Antibiosis of *Fusarium* spp. Applied and Environmental Microbiology 82, 1372–1382.

24. Rasheed, U., Ain, Q.U., Liu, B., 2024. Integration of Fe-MOF-laccase-magnetic biochar: From Rational Designing of a biocatalyst to aflatoxin B1 decontamination of peanut oil. Chemosphere 367, 143424.

25. Ren, Y., Yu, G., Shi, C., Liu, L., Guo, Q., Han, C. et al., 2022. Majorbio Cloud: A one-stop, comprehensive bioinformatic platform for multiomics analyses. Imeta 1, e12.

26. Sahithya, K., Mouli, T., Mercy Scorlet, T., 2022. Remediation potential of mushrooms and their spent substrate against environmental contaminants: An overview. Biocatalysis and Agricultural Biotechnology 42, 102323.

27. Shao, M.W., Chen, H.J., Huang, A.Q., Zheng, L., Li, C.j., Qin, D., et al., 2025. Modulation of rhizosphere microbiota by *Bacillus subtilis* R31 enhances long-term suppression of banana Fusarium wilt. iMetaOmics 2, e70006.

28. Ullah, M., Rahman, M., 2024. Adsorptive removal of toxic heavy metals from wastewater using water hyacinth and its biochar: A review. Heliyon 10, e36869.

29. Wang, H., Xu, M., Cai, X., Feng, T., Xu, W., 2020. Application of spent mushroom substrate suppresses Fusarium wilt in cucumber and alters the composition of the microbial community of the cucumber rhizosphere. European Journal of Soil Biology 101, 103245.

30. Xie, X., Huang, C., Cai, Z., Chen, Y., Dai, C., 2019. Targeted Acquisition of *Fusarium oxysporum* f. sp. *niveum* Toxin-Deficient Mutant and Its Effects on Watermelon Fusarium Wilt. Journal of Agricultural and Food Chemistry 67, 8536–8547.

31. Yang, H., Dai, H., Chen, Y., Wan, X., Li, F., Xu, Q., 2023. Efficient and simple simultaneous adsorption removal of multiple mycotoxins from environmental water. Separation and Purification Technology 317, 123888.

32. Zhang, J., Liu, Y., Zhang, N., Hu, B., Jin, T., Xu, H. et al., 2019. NRT1.1B is associated with root microbiota composition and nitrogen use in field-grown rice. Nature Biotechnology 37, 676–684.

33. Zhou, J., Wang, M., Sun, Y., Gu, Z., Wang, R., Saydin, A. et al., 2017. Nitrate Increased Cucumber Tolerance to Fusarium Wilt by Regulating Fungal Toxin Production and Distribution. Toxins 9, 100.

